# Cytokine interaction networks, not individual cytokines, drive anti-TNF response in Crohn’s disease

**DOI:** 10.64898/2026.08.11.744137

**Authors:** Marton Olbei, John P. Thomas, Yufan Liu, Sadek Malas, Dezso Modos, Nick Powell, Tamas Korcsmaros

## Abstract

Crohn’s disease (CD) is a chronic inflammatory condition of the gastrointestinal tract for which anti-tumour necrosis factor (anti-TNF) agents remain a first-line biologic therapy. However, remission rates are modest, and the mechanistic basis of non-response is poorly characterised. A common resistance mechanism is thought to emerge when alternative inflammatory cascades compensate for TNF inhibition, but the interactions underlying this rewiring have not been systematically characterised. We applied CytokineLink, our previously developed systems immunology framework, to single-cell RNA sequencing data from CD patients sampled before and after anti-TNF therapy. We reconstructed networks of interacting cytokines across samples stratified by treatment phase, response, and inflammation status, and identified condition-specific cytokine interactions and feedback loops, statistically validated against degree-matched random networks. We clustered the generated networks based on their inflammation, response, and treatment status. The pre-treatment inflamed non-responder network contained the largest set of unique interactions, organised around a connected module driven by IL17C targeting downstream TNF, IL6, IL1B, CXCL1/2/3/8, and CCL20. IL17C was produced by a population of non-ileal enteroendocrine cells, differentially abundant at baseline in non-responders. Gene set variation analysis in an independent cohort confirmed elevated non-responder module activity in colonic tissues of non-responders. Feedback loop analysis revealed that responder networks were characterised by persistent IL10 circuits sustained by macrophage populations and acquired tissue-remodelling interactions after therapy, whereas non-responders lost IL10 feedback loops post-treatment and gained TNF-containing motifs, including circuits signalling through the upstream activator TL1A. Our findings characterise the mechanism of anti-TNF non-response as a cytokine network, in which pre-existing epithelial-driven inflammatory modules and the failure to preserve regulatory feedback sustain TNF-independent inflammation in CD. By characterising cytokine interactions at the systems level, our approach moves beyond single-cytokine models of anti-TNF resistance to provide a mechanistic framework for understanding the biological basis of treatment failure in immune mediated diseases.

## Introduction

Crohn’s disease (CD) is a chronic inflammatory condition of the gastrointestinal tract, affecting up to 2.5 millions of people globally (Hracs et al., 2025; Kaplan, 2025). Biologic treatments targeting key cytokines have substantially improved outcomes for CD patients, in particular those targeting the proinflammatory cytokine tumour necrosis factor-alpha (TNF). Anti-TNF therapies such as infliximab and adalimumab are typically used as first-line biologic agents in clinical practice owing to their relative efficacy and cost-effectiveness. Recent studies have demonstrated that early top-down therapy with anti-TNF agents yields better 1 year outcomes in moderate-to-severe CD (Noor et al., 2025). However, reflecting the heterogeneity of CD, anti-TNF drugs still achieve relatively poor success rates compared to placebo, with net remission rates ranging from 17% to 29% (Kayal et al., 2023). Failure to respond to anti-TNF agents can be due to either primary non-response (failure to respond from the outset) or secondary loss of response (failure to respond over time) (Atreya, Neurath, & Siegmund, 2020), but the mechanistic basis for these remains poorly characterised. Understanding the mechanisms of anti-TNF resistance is needed to develop strategies that improve outcomes for CD patients.

While treatment failure can be caused by a variety of reasons, such as inadequate drug concentrations or the development of anti-drug antibodies, patients frequently exhibit resistance despite appropriate anti-TNF levels (Marsal et al., 2022). In such cases, resistance stems from underlying biology, such as alternative inflammatory cascades compensating for TNF inhibition. In anti-TNF resistant patients, studies have shown a shift toward other pro-inflammatory mediators that can sustain intestinal inflammation independently of TNF. One proposed pathway is IL23 receptor upregulation, and its downstream cytokine, IL17A, in anti-TNF non-responders compared to responders (Schmitt et al., 2019). However, while IL17A inhibition found success in other immune mediated inflammatory diseases (IMIDs) such as ankylosing spondylitis and psoriatic arthritis, anti-IL17A trials have failed in CD (Schmitt, Neurath, & Atreya, 2021). Similarly, targeting oncostatin-M (OSM), a cytokine highly expressed in IBD intestinal tissues which is associated with anti-TNF treatment resistance (West et al., 2017), has not yet yielded an effective therapy for CD.

The limited success of cytokine-targeting therapies reflects both the complexity and the redundancy of cytokine networks in CD. Although TNF is widely regarded as a central pro-inflammatory cytokine in CD (Souza, Caetano, Magalhães, & Castelucci, 2023), the disease is driven by a complex network of cytokines and immune cells (Vebr, Pomahačová, Sýkora, & Schwarz, 2023), of which TNF is only one, albeit critical node. Targeting TNF perturbs multiple downstream immune pathways, influencing the expression and activity of additional cytokines (Linares et al., 2022). The resulting cytokine milieu reflects a disease- and treatment-specific network state that emerges from these dynamic interactions, which can be computationally inferred with systems-level approaches (Jansen, Aschenbrenner, Uhlig, Coles, & Gaffney, 2022; Olbei et al., 2021).

To address this, we used single-cell RNA sequencing (scRNA-seq) data from site-matched intestinal biopsies from 16 biologic-naïve CD patients pre- and post- anti-TNF therapy <u>(Thomas</u> <u>et al. 2024)</u> and applied CytokineLink, our previously developed and validated systems immunology framework (Olbei et al., 2026), to deconvolute cytokine-cytokine interaction networks. We hypothesised that baseline responder versus non-responder networks exhibit distinct cytokine regulatory architectures, and that understanding how they regulate each other will be critical for explaining anti-TNF treatment resistance in CD. By resolving cytokine interactions on a systems level, our approach moves beyond single-cytokine descriptions of anti-TNF resistance, offering a novel mechanistic view of treatment failure.

## Results

### Treatment and response specific cytokine networks in Crohn’s disease

To understand the molecular mechanisms underlying anti-TNF treatment resistance in CD, we constructed cytokine networks using publicly available scRNA-seq data from the TAURUS study (Thomas et al., 2024). The integrated single-cell dataset was processed through the CytokineLink pipeline (Olbei et al., 2021; Olbei et al., 2026). In each resulting cytokine network, cytokines are represented as nodes and their interactions are predicted cytokine - cytokine regulatory interactions. Each interaction is cell type-specific, tracing the chain from: cytokine → receptor → signalling proteins → transcription factors → cytokine gene. We generated separate cytokine networks for each clinical state, categorised by treatment phase (pre- or post-treatment), responder status (responder- or non-responder based on clinical, endoscopic and/or histologic features as defined in the TAURUS study), and annotated biopsy sample inflammation status (inflamed or non-inflamed samples) (Figure 1A), yielding 8 clinical subgroups such as “pre-treatment inflamed non-responder” samples.

**Figure 1.**
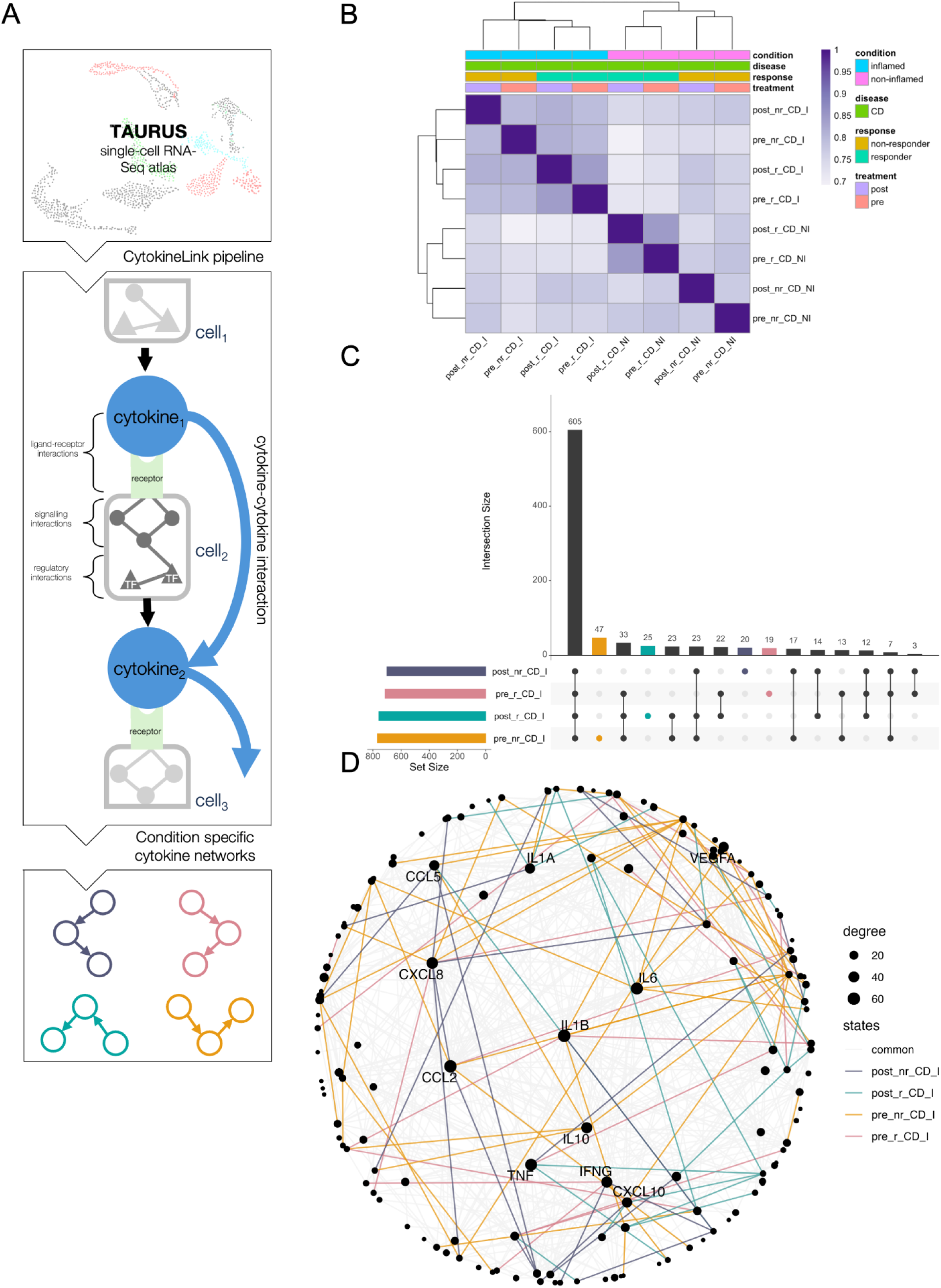
Treatment and response specific cytokine networks in Crohn’s disease. A: Schematic workflow of the study. Pre-processed scRNA-seq data published with Thomas et al., 2024 was downloaded from the project repository, and processed with our pipeline, that produces a separate cytokine network for each condition. B: Heatmap showing the pairwise Jaccard-indexes of the generated networks. C: UpSet plot displaying the number of shared interactions between the inflamed networks. D: Network visualization of all interactions within the inflamed networks, highlighting high-degree (high number of cytokine interactors) cytokines. Betweenness centrality layout positions the most highly-connected cytokines at the core. Labels: pre_nr_CD_I = pre-treatment non-responder Crohn’s Disease inflamed, pre_nr_CD_NI = pre-treatment non-responder Crohn’s Disease non-inflamed, pre_r_CD_I = pre-treatment responder Crohn’s Disease inflamed, pre_r_CD_NI = pre-treatment non-responder Crohn’s Disease non-inflamed, post_nr_CD_I = post-treatment non-responder Crohn’s Disease inflamed, post_nr_CD_NI = post-treatment non-responder Crohn’s Disease non-inflamed, post_r_CD_I = post-treatment responder Crohn’s Disease inflamed, post_r_CD_NI = post-treatment responder Crohn’s Disease non-inflamed

We calculated the pairwise Jaccard indices between all 8 cytokine networks for each clinical subgroup, and hierarchical clustering of these data (Figure 1B) confirmed clustering according to inflammation, response, and treatment status. All networks shared Jaccard indices > 0.7, indicating substantial overlap in cytokine interactions across conditions. The UpSet plot on Figure 1C further illustrates the substantial overlap of interactions among cytokine networks within inflamed CD subgroups, alongside a smaller subset of state-specific interactions. The pre-treatment non-responder network contained the largest number of unique cytokine interactions (47).

### A unique cytokine module driven by IL17C characterises baseline non-responders

We next honed in on the 47 cytokine interactions exclusive to pre-treatment, inflamed samples in CD non-responders to anti-TNF therapy, which was the network with the largest number of unique cytokine interactions. 31 out of the 47 cytokine interactions belonged to a large connected subgraph, where the cytokines were connected primarily by IL17C and CCL16 (Figure 2A). We termed this the non-responder module. IL17C signaled 11 other cytokines within this module, while CCL16 sent 8 interactions, making up 51% of interactions in the subgraph. The most targeted cytokine was IL6, receiving 5 interactions from other nodes in the module (Figure 2B).

**Figure 2.**
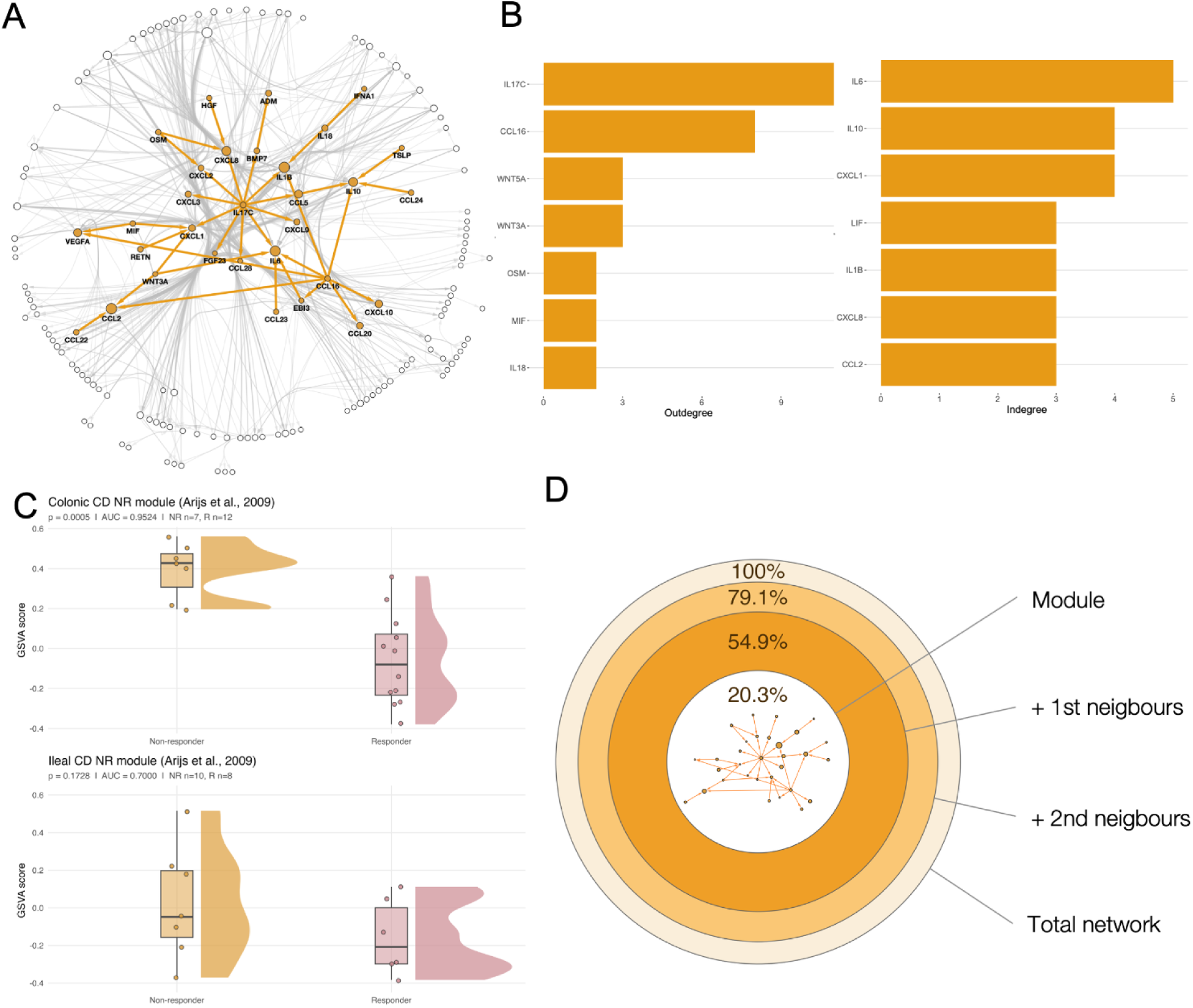
A unique IL17C-driven cytokine module characterises baseline non-responders. A: Cytokine interaction network of pre-treatment non-responders, with interactions exclusive to this condition highlighted in orange. B: Bar plots show out-degree and in-degree of cytokines within the module. C: Non-responder module activity in an independent cohort (Arijs et al., 2009), comparing anti-TNF responders and non-responders. Module activity (GSVA score) is increased in colonic non-responders but does not differ in the ileum. GSVA, gene set variation analysis. D: Reach of the IL17C module through the pre-treatment non-responder network. Concentric rings show the cumulative percentage of the network encompassed at successive steps outward from the module (module → first neighbours → second neighbours → whole network), with ring area proportional to that percentage. The module together with its first-neighbour targets (including TNF, IL6, IL1B, CXCL1/2/3/8 and CCL20) accounts for 54.9% of the network.

To validate the relevance of the non-responder module in independent data, we performed gene set variation analysis (GSVA) of the module’s constituent genes in the deposited microarray dataset of Arijs et al., 2009 (Arijs et al., 2009), in which the authors measured longitudinal gene expression differences between responders and non-responders to anti-TNF therapy, split by anatomical location. GSVA analysis of the full non-responder module revealed strong discriminatory performance in pre-treatment colonic biopsies for distinguishing anti-TNF responders from non-responders (Wilcoxon p = 0.0005, AUC = 0.95 (95% CI 0.829–1.000), n = 19 colonic CD patients). However, this was not observed in ileal biopsies (p = 0.1728, AUC = 0.700 (95% CI 0.407–0.922), n = 18 ileal patients), indicating that the module’s discriminatory signal is colon-specific (Figure 2C). This colonic restriction is consistent with a non-ileal epithelial origin for the module, which we localise to a specific epithelial population below.

Based on our cytokine network model, the non-responder module has broad downstream consequences. Through their first-neighbour targets, which include pro-inflammatory cytokines and chemokines such as TNF, IL6, IL1B, CXCL1, CXCL2, CXCL3, CXCL8, and CCL20, the module impacts 54.9% of the entire cytokine network in pre-treatment non-responders (Figure 2D). The module could therefore amplify a broad inflammatory programme.

Each cytokine-cytokine interaction in our networks is supported by one or more cell type-specific signalling chain. A sender cell type expressing the upstream cytokine, and a receiver cell type expressing its receptor, the downstream signalling proteins and transcription factors, and the target cytokine gene. Every network therefore carries a cell type layer, in which each cell type can be described by the set of cytokines it sends and receives, that is, by its cytokine signalling neighbourhood. We used this layer to study which cell types account for the interactions specific to pre-treatment non-responders. We collated the cell-cytokine interactions underlying each edge and compared the cytokine signalling neighbourhood of every cell type between pre-treatment responders and non-responders by rewiring analysis with DyNet. A highly rewired cell type is one that sends or receives a different set of cytokines between the two conditions, in number, identity, or both. Top-rewired cell types overlapped significantly with differentially abundant cells (hypergeometric test, *P* = 0.002) (Figure 3A), indicating that differentially abundant cell types receive and secrete a more variable set of cytokines, which in turn affect downstream cytokines. These cell types therefore differ not only in abundance but also in cytokine signalling function. In the non-responder module, we noted that 46% of edges (cytokine-cytokine interactions) involved at least one differentially abundant cell type.

**Figure 3.**
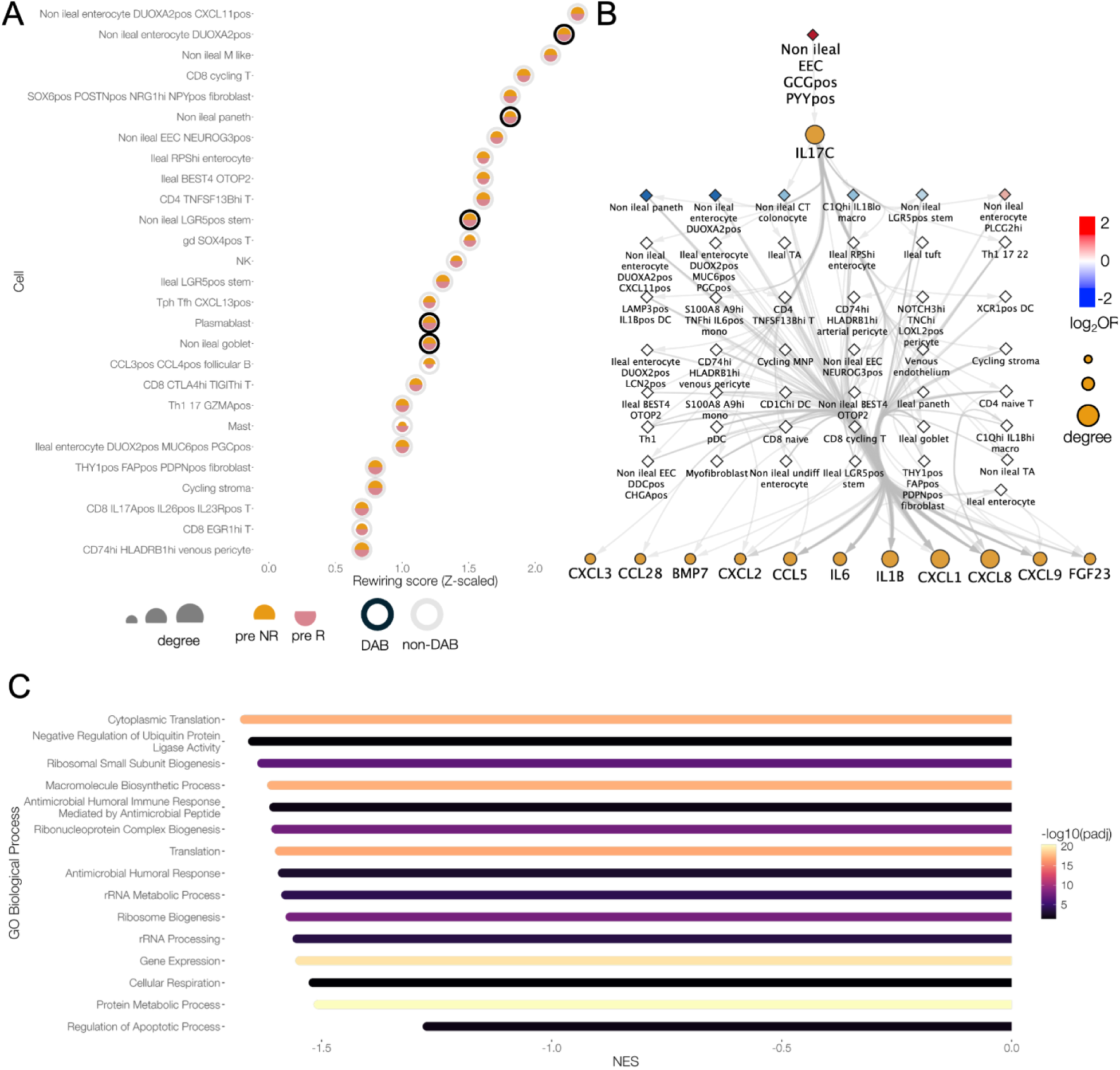
Differentially abundant epithelial cells drive the IL17C module. A: Cell rewiring between pre-treatment responder and non-responder samples, with cell types ranked by rewiring score (Z-scaled). Top-rewired cells (Z > 1) are enriched for differentially abundant (DAB) cells (hypergeometric P = 0.002). B: IL17C is produced by one of the most differentially abundant cell types between pre-treatment responder groups, non-ileal GCG⁺ PYY⁺ enteroendocrine cells. Node fill of cell type nodes indicates the log2 odds ratio of differential abundance (non-responders versus responders) and node size indicates degree. C: Gene set enrichment analysis of the non-ileal GCG⁺ PYY⁺ enteroendocrine cell population between non-responders and responders, highlighting the negative enrichment of antimicrobial defense pathways.

We next investigated which cell populations drive the non-responder module. IL17C was produced by one of the most differentially abundant cell type expanded in pre-treatment non-responders, a non-ileal GCG⁺ PYY⁺ enteroendocrine cell (EEC) population (OR = 7.2, Benjamini-Hochberg *P* = 0.001 as reported in Thomas et al. 2024, Supplementary Table 6) (Figure 3B). Gene set enrichment analysis (GSEA) of this non-ileal GCG⁺ PYY⁺ EEC population between pre-treatment non-responders versus responders revealed significant negative enrichment in gene ontology terms related to antimicrobial defense pathways in non-responders (Figure 3C), involving genes involved in microbial pattern recognition such as DMBT1 (gp-340) and LGALS3 (Galectin-3) (Díaz-Alvarez & Ortega, 2017; End et al., 2009) and effectors such as REG1B (Qi et al., 2025). This suggests that one of the most expanded epithelial cell types at baseline in non-responders may be functionally impaired in its ability to mount protective responses to microbial stimuli.

IL17C has been shown to be induced by microbial stimuli at mucosal surfaces, particularly in enteroendocrine and goblet cells (Swedik, Madola, & Levine, 2021). Supporting this, data from the IBD Model Bank (Gonzalez-Acera et al., 2025) revealed that overtly bacteria-driven murine colitis models (e.g. *Helicobacter hepaticus* and *Citrobacter rodentium* infection models) exhibit stronger induction of IL17C than other colitis models (Figure S2). Together, these observations suggest that impaired antimicrobial defense capabilities of the expanded GCG⁺ PYY⁺ EEC population in inflamed CD tissues of non-responders, may predispose to augmented microbial-driven IL17C secretion, although this requires further experimental validation .

Collectively, the module’s broad first-neighbour reach and its origin in an antimicrobially compromised epithelial population suggest a mechanism by which baseline inflammation could resist TNF blockade. The IL17C signal diminishes after treatment in non-responders; however, its pre-treatment presence may contribute to primary non-response, making IL17C both a baseline biomarker and a mechanistic driver of resistance. Further mechanistic studies are needed to confirm this causal link.

### Responder specific feedback loops center around IL10, while non-response specific feedback loops center around TNF

To better understand the role of cytokine interactions in their respective cytokine networks, we enumerated all three-node feedback loops (A → B → C → A) present in each of the generated cytokine networks. These loops represent network motifs, recurring patterns of interactions, that serve as the basic building blocks of complex systems. Feedback loops are built from the same underlying directed interactions that comprise the cytokine networks described above, and most feedback loops are shared across response groups and treatment states. A substantial backbone of motifs is present in every condition, with 144 motifs identified in all four states, involving many of the same hub cytokines (IL1B, CCL2, IL10, TNF, IL6, CXCL10). This conserved backbone suggests that the core regulatory architecture of the network is preserved regardless of treatment outcome, and that response biology is therefore likely captured by the smaller set of circuits that differ between conditions, which we studied next.

We identified feedback loops exclusive to responders or non-responders and classified each as persistent (present in both pre- and post-treatment networks in a response group) or emergent (appearing only post-treatment). Additionally, we identified feedback loops exclusive to baseline responders or non-responders, which represent circuits lost after therapy (Figure 4A). We focussed our analysis on the few motifs that were different between conditions, to see if differences in treatment response may be encoded in these differentiating feedback loops. To increase our confidence in the relevance of these condition-specific motifs, we established their prevalence in 10,000 degree-matched randomised networks where 40 out of 50 motifs achieved an adjusted p-value ≤ 0.05 (Figure 4B). The identified motifs form connected subgraphs in their respective states (Figure 4C)

**Figure 4.**
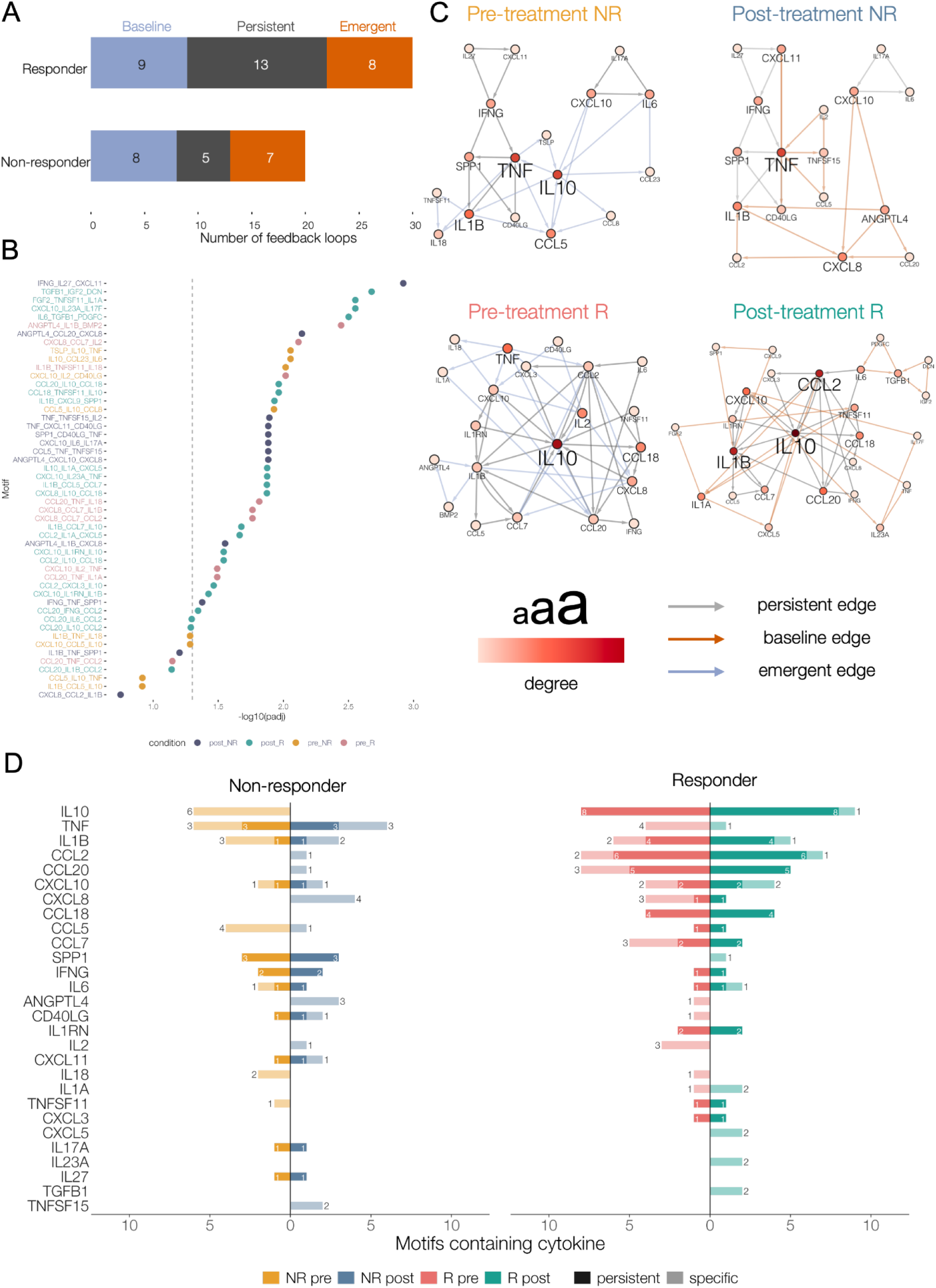
Feedback loops are uniquely present in post-treatment non-responders and responders. A: Number of response group specific motifs at baseline, post-treatment, and persistent through treatment. B: Significant motifs following based on their occurrence in 10,000 degree-matched random networks. C: The response group specific motifs form connected subnetworks. Fill colour and label size correspond to degree, edge colour corresponds to the persistent, baseline or emerging status of the edges. D: Number of occurrences per cytokine, response group and treatment state.

In total, we identified 21 responder-specific and 12 non-responder-specific post-treatment feedback loops. In responders, 13 of 21 motifs were persistent and 8 were emergent. In non-responders, 5 of 12 were persistent and 7 were emergent. At baseline, 8 feedback loops were exclusive to pre-treatment non-responders and 9 were exclusive to pre-treatment responders.

In pre-treatment responders 4 out of 9 feedback loops involved TNF. These TNF-containing loops were absent post-treatment, consistent with effective TNF blockade. The post-treatment responder feedback loop network converged on IL10, which had the highest motif in-degree with 9 of 21 responder-specific motifs terminating on it (Figure 4D). Of these, 8 were persistent (present both before and after treatment) indicating that the specific IL10 feedback in responders is a stable feature of the network, maintained through therapy. The persistent IL10-containing motifs involved monocyte- and macrophage-recruiting chemokines such as CCL18, CCL2 (MCP-1), CCL7 (MCP-3), and CCL20 (MIP-3α). The most prolific source cell types contributing the unique edges of responder-specific feedback loops were macrophages or monocytes (Figure S3). As for emergent responder specific motifs, only one involved IL10. Two motifs involved tissue remodelling factors: IL6 - TGFB1 - PDGFC and TGFB1 - IGF2 - DCN, comprising growth factors and extracellular matrix components. These features suggest that response is characterised not just the removal of TNF-driven inflammation but the presence of a stable IL10 regulatory programme and the acquisition of tissue-remodelling circuits, a signature consistent with a shift from active inflammation toward resolution and tissue repair.

In non-responders, 8 feedback loops were exclusive to pre-treatment non-responders. Six of these contained IL10, with CCL5 serving as a frequent intermediary node (present in 4 of the 6 IL10-containing motifs). However, no IL10-containing motifs were detected in the post-treatment non-responder network, indicating that these IL10 feedback circuits are present at baseline but are lost following anti-TNF therapy in non-responders.

The post-treatment non-responder feedback loop network instead centered on TNF, which had the highest motif in-degree with 6 of 12 motifs involving it. Three of these TNF centered motifs were persistent, and three were emergent. Two of the emergent loops connected TNF through TL1A, an upstream activator of TNF whose activity is not suppressed by anti-TNF therapy <u>(Jin</u> <u>et al. 2013)</u>. There were two persistent TNF-free motifs, CXCL10 - IL6 - IL17A and IFNG - IL27 - CXCL11, and four emergent TNF-free feedback loops, mainly involving ANGPTL4 and CXCL8. Non-responders show a mirrored image of the responder pattern. The IL10 regulatory circuits present at baseline are lost, while TNF becomes most prominent in the post-treatment network through both persistent and newly emergent feedback loops, including TL1A-driven motifs.

Several cytokines participated in feedback loops in both conditions, including CXCL10, IL1B, IL6, CCL2, CXCL8, and CCL20, but had different positions within the motif networks. In responders, CXCL10 participated in circuits with IL1RN and IL10, in non-responders, CXCL10 participated in the persistent CXCL10 - IL6 - IL17A loop. IL1B appeared in IL10 motifs in responders, and CXCL8 motifs in non-responders (Figure 4D). Taken together, these findings show that the structure of the cytokine network distinguishes response from non-response to anti-TNF therapy in Crohn’s disease: responders retain IL10-centred feedbacks, and acquire tissue repair-associated motifs, whereas non-responders after therapy lose IL10 regulation and maintain or acquire TNF-containing and TNF-independent inflammatory feedback loops.

## Discussion

A major challenge in Crohn’s disease and IBD is understanding why some patients fail to respond to anti-TNF therapy. Here, we show that anti-TNF non-response is encoded not by the behaviour of individual cytokines, but by the topology and rewiring of the broader cytokine network. Using Crohn’s disease scRNA-seq data from the TAURUS study, with samples collected at baseline and following anti-TNF treatment in responders and non-responders, we reconstructed cytokine networks across inflammatory and response states. These networks separated according to inflammation, treatment, and response status and revealed distinct state-specific interactions and network modules. At baseline, non-responders were characterised by a highly connected, epithelial IL17C-driven inflammatory module. Following treatment, they exhibited loss of regulatory IL10 feedback and emergence of additional TNF- and TL1A-containing circuits that sustained inflammation independently of TNF inhibition. These findings suggest that anti-TNF resistance is a systems-level property of the inflammatory network, in which the pre-existing network structure and subsequent rewiring enable inflammation to persist despite therapeutic targeting of a dominant cytokine.

Previous studies have identified individual cytokines, such as OSM (West et al., 2017), IL23/IL17A (Schmitt et al., 2019), and various transcriptomic signatures, that are differentially expressed in anti-TNF non-responders. While these approaches can identify what is different in non-responders, they do not fully explain why these cytokines are elevated, or how the non-responder inflammatory state emerges from specific cellular differences. Our cytokine network approach addresses this gap by resolving which cytokines distinguish non-responders, and the regulatory relationships between them. This information is hidden from differential expression or gene-set-level analyses alone.

In this study, we found the largest connected condition-specific cytokine module in pre-treatment, non-responding inflamed Crohn’s disease samples. This network module was organised around epithelial-derived IL17C, targeting pro-inflammatory cytokines and chemokines, including IL6, IL1B, CXCL1/2/3, CXCL8 and CCL20. Epithelial populations such as enteroendocrine cells produce IL17C, which drives inflammation by inducing multiple cytokines and chemokines we found targeted by IL17C in our study, such as CXCL1/2/3, CXCL8 (IL8), CCL20, IL1B, IL6 and TNF (Nies & Panzer, 2020). The levels of IL17C have also previously been found to be elevated in CD patients with active disease (M Friedrich, Diegelmann, Schauber, Auernhammer, & Brand, 2015). We found that cell types that were found differentially abundant at baseline between responder groups are highly rewired in the cytokine network, and contribute to this unique IL17C/CCL16 cytokine signalling module in pre-treatment non-responders. While the TAURUS study observed overall lower epithelial cell frequency at baseline in CD non-responder patients, the GCG⁺PYY⁺ EEC population was specifically expanded, suggesting enrichment of this cell type concurrent with broader epithelial barrier disruption. Independent validation by GSVA in the Arijs et al. 2009 microarray data confirmed that the module’s constituents are significantly elevated in non-responders (AUC = 0.95), specifically in colonic tissues, but not ileal ones, consistent with the colonic epithelium origin of the IL17C producing differentially abundant cell type.

IL17C is produced by epithelial-mucosal populations (Swedik et al., 2021), induced by bacterial stimuli (particularly in enteroendocrine cells), dependent on IL17A/TNF signalling, and elevated in active CD. Given the downregulated microbial-defence pathways we identified, we hypothesised that this differentially abundant cell type amplifies an IL17C-driven inflammatory cascade that compensates for TNF blockade. On a cytokine interaction level this functions by sustaining neutrophil-recruiting chemokines (CXCL1/2/3/8) and other inflammatory programs through IL1B / IL6, that have been shown to be enriched in anti-TNF non-responders (Aschenbrenner et al., 2021; Matthias Friedrich et al., 2021). These observations raise the possibility that an epithelial IL17C signature may help maintain a TNF-independent inflammatory circuit in Crohn’s disease, and therefore may contribute to primary non-response.

In previous studies, serum level differences between anti-TNF responders and non-responders established higher levels of TGFB1 and MMP-9 with lower sCD14 at baseline, that may indicate greater barrier damage and altered microbial sensing already before therapy begins in these patients (Coufal et al., 2023). In the Thomas et al. 2024 dataset we have found that ileal and non-ileal epithelial cell populations in general both were negatively enriched for direct microbial defense pathways between responders and non-responders, indicating that the latter group may be suppressed in their ability to repel the microbiome. The IBD Model Bank data provided further support for a microbiome-driven origin of the IL17C signal we observed (Gonzalez-Acera et al., 2025). IL17C induction in bacterially-driven, but not barrier-damage colitis models supports the hypothesis that that microbial engagement (potentially amplified by the impaired antimicrobial defenses of the GCG⁺ PYY⁺ EEC population) may activate the IL17C signal observed at baseline in non-responders. Whether this reflects a pre-existing dysbiotic state, an inability to contain microbiome translocation, or altered pattern recognition in this cell type remains to be determined. The lack of responder-stratified studies using direct barrier function readouts in Crohn’s disease accentuates the need for future work in this area, including targeted receptor mapping, organoid perturbation studies, and prospective clinical correlation to make a causal link, and evaluate IL17C as a key cytokine of anti-TNF therapy resistance.

The condition-specific motifs analysed in our work represent a minority of the total feedback structure, as the majority is shared across response groups and treatment states. We focussed on the differentiating subset because it is where response biology may be encoded. Here, our feedback loop analysis showed that anti-TNF therapy rewires higher-order network structures rather than individual nodes, and that the direction of this rewiring differs between response groups. In responders, baseline TNF-containing feedback loops were lost after treatment, consistent with successful TNF blockade, while IL10-centred feedback loops persisted through therapy. This agrees with reports that successful anti-TNF therapy depends on an IL10-dependent macrophage programme <u>(Koelink et al. 2020)</u>, and with our finding that the C1Qhi IL1Blo macrophage population involved in sustaining these loops was itself differentially abundant in Thomas et al. (OR=0.46) (Figure S3). The responder-specific motifs that emerged also involved tissue-remodelling factors, (TGFB1, IGF2, PDGFC, DCN), suggesting that response involves both suppression of inflammation and induction of repair programmes.

In non-responders, the pattern was the inverse. The majority of pre-treatment-exclusive feedback loops contained IL10, often connected through CCL5 (itself a direct IL17C target), and these circuits were lost after treatment. Non-responders instead gained TNF-containing and TNF-independent pro-inflammatory motifs, including new circuits signalling through TL1A (TNFSF15), which acts upstream of TNF and is insensitive to anti-TNF blockade <u>(Jin et al. 2013;</u> <u>Parigi and Cominelli 2026)</u>. This persistent incoming TNF signalling matches reports of persistent TNF expression in non-responder monocytes despite therapy <u>(Gaiani et al. 2020)</u>. The remaining persistent TNF-independent loops are pre-existing Th17 and Th1 circuits that anti-TNF therapy may not disrupt. IL6, the most targeted node in the pre-treatment non-responder module (Figure 2A), along with the other downstream effectors of the module, becomes a constituent node of both persistent and emergent post-treatment loops in non-responders. This implies that the baseline IL17C module may help establish the compensatory circuits that sustain inflammation after therapy. Overall, our results suggest that anti-TNF therapy disrupts a pre-existing regulatory programme in patients who fail to respond, while leaving pro-inflammatory circuits intact.

Our study has limitations. The cytokine networks generated here are computationally inferred from transcriptomic data, and as such represent predicted regulatory interactions rather than experimentally validated signalling events. While the pipeline we employ has been benchmarked against experimental data, including IBD patient derived organoids and their co-culture with immune cells (Olbei et al., 2026), the cytokine network should be interpreted as a hypothesis-generating framework rather than a definitive map of active cytokine signalling. As a corollary, scRNA-seq captures transcriptional state and not protein secretion, and cytokine biology is known to be heavily post-transcriptionally regulated. The disconnect between mRNA expression and secreted protein levels is a known limitation of transcriptomics-based inference of cytokine activity (Schäfer, Dimitrov, Villablanca, & Saez-Rodriguez, 2024). The relatively modest cohort size of the TAURUS study, while longitudinal and carefully phenotyped, means that the group-aggregated networks may not fully capture the heterogeneity of treatment response across the broader CD patient population. Additionally, while we observe that the GCG⁺ PYY⁺ enteroendocrine cell population is negatively enriched for antimicrobial defense pathways and implicated as the primary source of IL17C in non-responders, the causal link between impaired microbial defense, microbiome engagement, and elevated IL17C secretion remains hypothetical in the absence of direct validation. The IBD Model Bank data we use to support the microbiome-IL17C connection, while suggestive, derives from animal models that do not fully recapitulate human CD biology, as important differences exist between human and murine immune systems (Mestas & Hughes, 2004), and cross-species inference should be treated with caution. The GSVA validation in the Arijs et al. cohort, while yielding statistically significant AUC values for the full module, was performed in a relatively small cohort. While the network-predicted module has a detectable transcriptional footprint in independent data, validation in larger cohorts, with matched protein-level measurements would be important to confirm the clinical applicability of these findings. The feedback loop analysis, while statistically validated against degree-matched random networks, is limited by the same inference caveats as the broader network, where the motifs represent predicted regulatory circuits, not experimentally confirmed signalling events.

Future work should prioritise experimental validation of the key network predictions, such as the proposed IL17C-driven inflammatory cascade in primary non-responders. Organoid perturbation studies using patient-derived intestinal organoids from responders and non-responders, in which the GCG⁺ PYY⁺ EEC population can be challenged with microbial stimuli and IL17C secretion directly measured, would provide an experimental system to test the central hypothesis. The loss of IL10 circuits in non-responders warrants experimental investigation, establishing whether IL10 feedback stability can be measured as a predictor of treatment response, and if interventions that stabilise IL10 signalling could improve outcomes in patients at risk of non-response. Whether the IL17C epithelial signature and the TNF feedback loops represent independent mechanisms of resistance or interact within a shared inflammatory program remains an open and important question for future studies.

In conclusion, the reconstruction of cytokine interaction networks points out that anti-TNF non-response in CD is a systems-level phenomenon and provides a route towards mechanistically informed patient stratification. Our analysis identified a pre-existing, epithelial IL17C-driven module that induces downstream pro-inflammatory cytokines that TNF neutralisation does not reach, while the regulatory IL10 feedback that helps stabilise the responder network is not maintained in patients who fail therapy. Non-response therefore can be considered as an inflammatory cytokine network structure that anti-TNF treatment leaves largely intact, rather than the failure to suppress one cytokine. For CD, this view points to several testable directions for improving care: an IL17C driven module detectable before treatment that could help identify patients at risk of primary non-response, upstream nodes such as TL1A that remain active despite TNF blockade and are currently undergoing clinical trials, and the stability of macrophage-supported IL10 feedback as a candidate predictor of durable response.

More broadly, the framework applied here is not specific to Crohn’s disease or to anti-TNF therapy. Any immune-mediated inflammatory disorder (IMID) in which responders and non-responders can be sampled at single-cell resolution can be analysed with the presented methodology. Because redundancy and compensation between cytokines can make the efficacy of single-target strategies difficult to predict, mapping how a network is wired could be an effective option for understanding treatment resistance across inflammatory diseases, and for guiding patient selection and combination therapy in IMIDs.

## Methods

### Generation of cytokine networks

Integrated, pre-processed h5ad files containing the individual major cell types (i.e. “myeloid”, “cd4” etc.) were downloaded from the TAURUS Zenodo repository (https://zenodo.org/records/14007626), and read into R (version 4.5.2) as Seurat (version 5.4.0) (Hao et al., 2024) objects using the schard package (version 1.0.0). Cell types were combined into objects based on the following metadata: disease (i.e. “CD”, “Healthy”), treatment (“pre-treatment”, ”post-treatment”), inflammation (“inflamed”, ”non-inflamed”) and remission (“responder”, “non-responder”) status. Patient specific networks were generated in an identical manner, but data was further split according to sample identifiers (i.e. “R_CD13”). The resulting Seurat objects (i.e. “pre_responder_CD_inflamed”) were processed with our CytokineLink pipeline to generate cytokine networks, using the “final_analysis” cell type identity provided by the original authors (Olbei et al., 2026). Briefly, the pipeline uses NicheNet (version 2.2.1.1) (Browaeys, Saelens, & Saeys, 2020) to establish active cytokine ligands, and their putative cytokine target genes in all cell types included in the dataset, and generates state-specific cytokine interaction networks as its output. For more information on the pipeline please refer to the Methods section in our previous work (Olbei et al., 2026).

### Analysis of cytokine networks

#### Network similarity and hierarchical clustering

The Jaccard-index of each cytokine network pair was calculated in R with a custom function. Hierarchical clustering of the cytokine networks was established with the pheatmap (version 1.0.13) library.

#### Network analysis

The generated state-specific cytokine networks were analysed in R, centrality measures were calculated using the tidygraph (version 1.3.1) and ggraph (version 2.2.2) packages.

#### Network rewiring

The DyNet (version 1.0) library was used to calculate the rewiring values between the network states Cytoscape (version 3.9.1), using multiple comparison mode, results were exported to .csv files and further processed in R for visualisation. Cell types were considered highly rewired when their Z-scaled rewiring values were > 1. The significance of the overlap with differentially abundant cell types was calculated with the base R *phyper* function.

#### Network motifs and statistical testing

Feedback loops were established using the ISMAGS Cytoscape app (version 1.0.5). Briefly, a template A → B, B → C, C → A triangle motif was created as a new network in Cytoscape, and this motif was mapped against the four loaded condition-specific Crohn’s disease networks (pre-/post treatment, responder/non-responder). The resulting list of matching motifs was exported, and further processed in R, establishing feedback loops shared between and distinct to each condition.

To establish the statistical significance of the generated condition specific motifs, we generated 10,000 directed, degree preserving random networks (i.e. edges were swapped in a way that keeps in- and outdegree values) from both the responder and non-responder specific networks, using the sample_degseq function in R igraph (version 2.2.1). In each iteration, in each random network, the occurrence of each tested motif was recorded with a 1 or its absence with a 0. For each motif, the fraction of occurrences from the 10000 simulations was used as a Monte Carlo p-value.

#### GSVA analysis

To assess whether the constituents of non-responder module can discriminate responders and non-responders in independent datasets, gene set variation analysis (GSVA) was performed using the GSVA R (version 2.4.4) package on the deposited microarray dataset of Arijs et al., 2009 (GSE16879). Data was retrieved from GEO using the GEOquery (version 2.78.0) (Davis & Meltzer, 2007) library. GSVA scores were compared between pre-treatment responders and non-responder samples using Wilcoxon rank-sum test. Performance was quantified by AUC using the pROC library (version 1.19.0). Analysis was performed separately for both colonic and ileal samples.

#### Gene set enrichment analysis

Differentially expressed genes between pre-treatment non-responders and responders were identified separately within ileal and non-ileal epithelial cell populations (and within the non-ileal GCG⁺PYY⁺ enteroendocrine subset specifically) using the FindMarkers function in Seurat. Genes were ranked by log2 fold-change and tested for enrichment of Gene Ontology Biological Process terms using fast gene set enrichment analysis (fgsea, version 1.36.2) (Korotkevich et al., 2016). Pathways with a Benjamini-Hochberg adjusted p-value ≤ 0.05 were considered significantly enriched.

#### IBD Model Bank comparison

Differential expression of Il17c across murine colitis models was obtained directly from the IBD Model Bank resource (Gonzalez-Acera et al., 2025), which provides pre-computed log2 fold-changes and Benjamini-Hochberg adjusted p-values for each model relative to its respective control. No additional statistical testing was performed locally, models were ranked and visualised by their reported adjusted p-value and fold-change.

## Acknowledgements

We thank the past and present members of the Korcsmaros and Powell groups for the advice and suggestions regarding our work.

## Data availability

All codes to reproduce the results in this manuscript can be found in the project github repository https://github.com/korcsmarosgroup/CD-antiTNF-cytokine-networks and zenodo (10.5281/zenodo.21805981). The generated networks are available on the NDEx network repository.

## Supplementary materials

**Figure S1:**
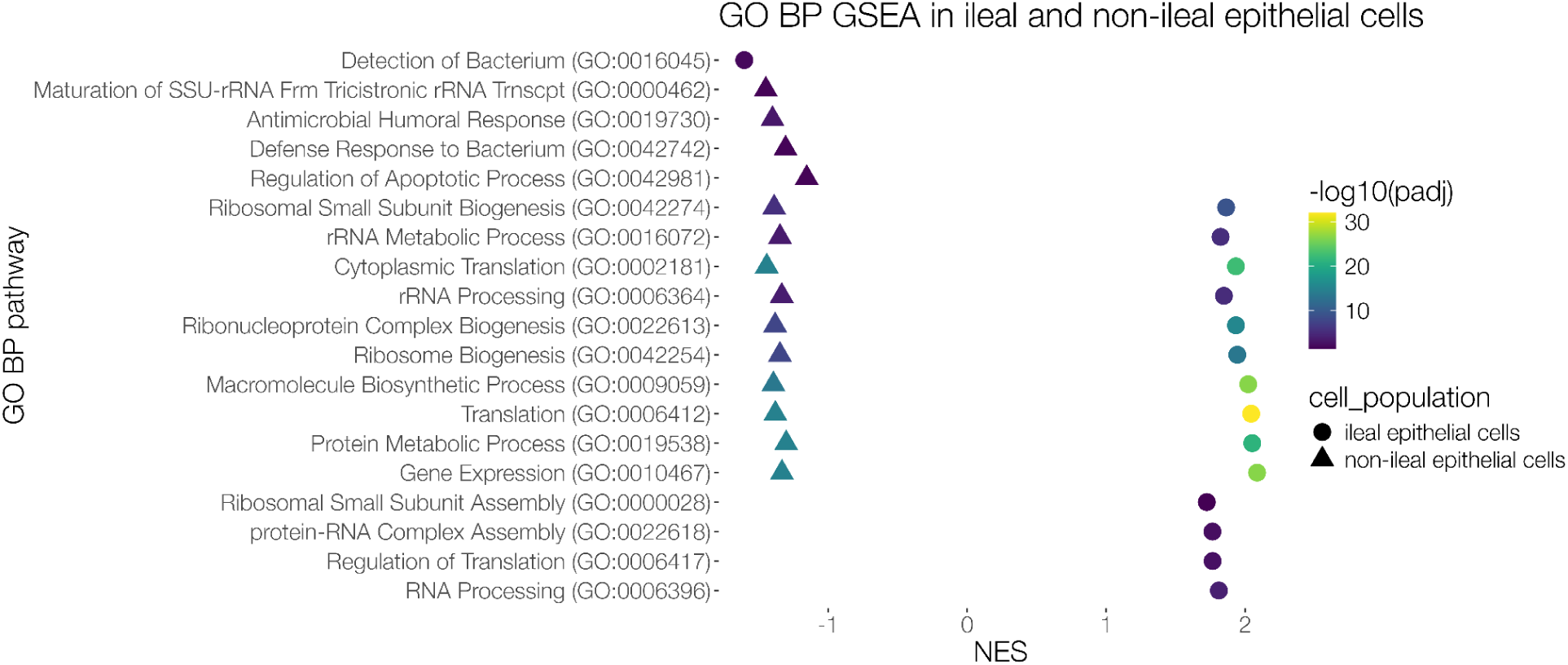
Gene Ontology pathway enrichment in ileal and non-ileal epithelial cells. Ileal and non-ileal epithelial pathways are both negatively enriched for microbial defense pathways, highlighted in red.

**Figure S2:**
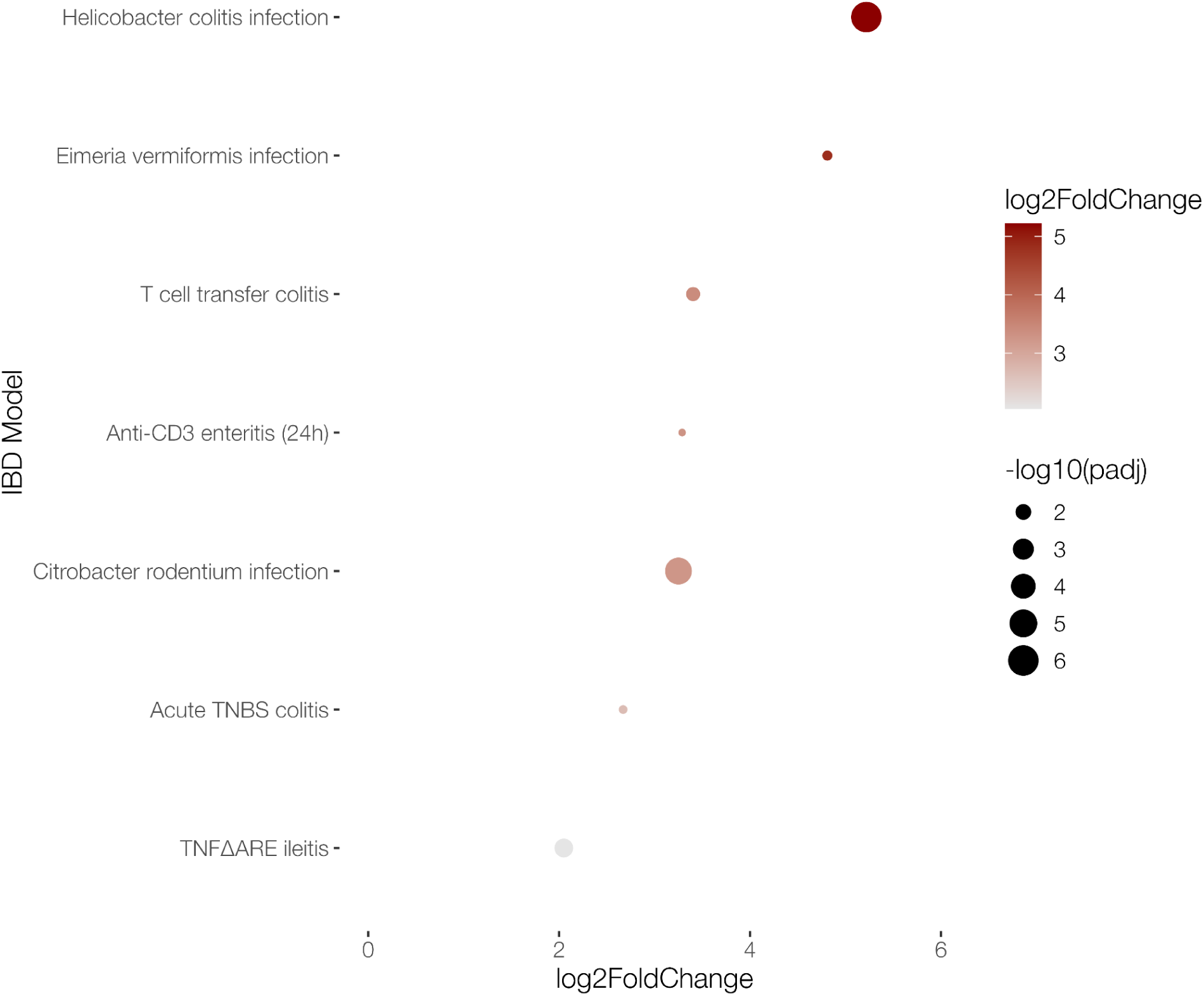
**IL17C response in murine models of inflammation**

**Figure S3:**
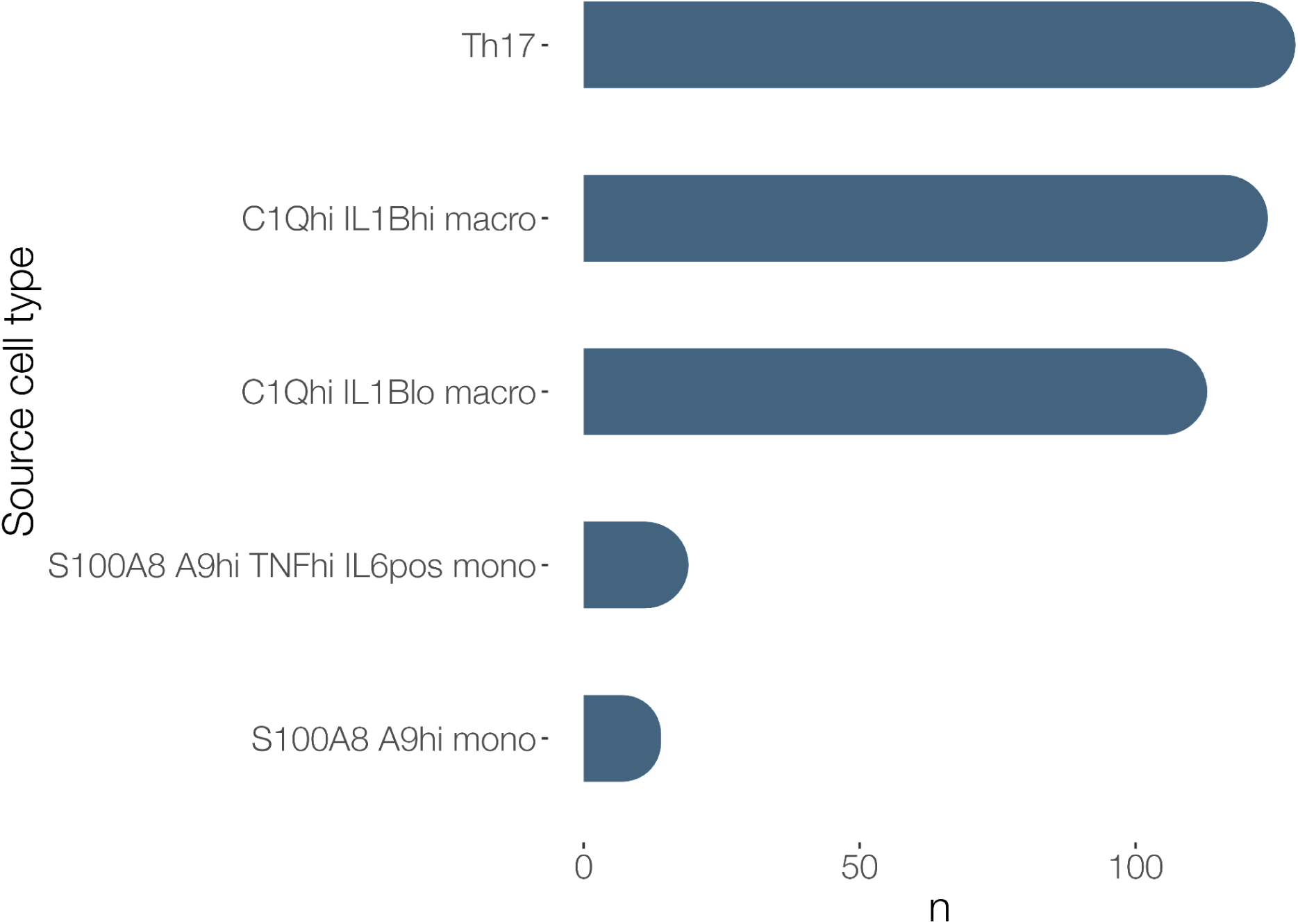
**Source cell types of the unique edges in responder-specific feedback loops centered around IL10. n = number of interactions**

## Bibliography

Arijs, I., De Hertogh, G., Lemaire, K., Quintens, R., Van Lommel, L., Van Steen, K., Leemans, P., et al. (2009). Mucosal gene expression of antimicrobial peptides in inflammatory bowel disease before and after first infliximab treatment. Plos One, 4(11), e7984.

Aschenbrenner, D., Quaranta, M., Banerjee, S., Ilott, N., Jansen, J., Steere, B., Chen, Y.-H., et al. (2021). Deconvolution of monocyte responses in inflammatory bowel disease reveals an IL-1 cytokine network that regulates IL-23 in genetic and acquired IL-10 resistance. Gut, 70(6), 1023–1036.

Atreya, R., Neurath, M. F., & Siegmund, B. (2020). Personalizing Treatment in IBD: Hype or Reality in 2020? Can We Predict Response to Anti-TNF? Frontiers in medicine, 7, 517.

Browaeys, R., Saelens, W., & Saeys, Y. (2020). NicheNet: modeling intercellular communication by linking ligands to target genes. Nature Methods, 17(2), 159–162.

Coufal, S., Kverka, M., Kreisinger, J., Thon, T., Rob, F., Kolar, M., Reiss, Z., et al. (2023). Serum TGF-β1 and CD14 Predicts Response to Anti-TNF-α Therapy in IBD. Journal of immunology research, 2023, 1535484.

Davis, S., & Meltzer, P. S. (2007). GEOquery: a bridge between the Gene Expression Omnibus (GEO) and BioConductor. Bioinformatics, 23(14), 1846–1847.

Díaz-Alvarez, L., & Ortega, E. (2017). The Many Roles of Galectin-3, a Multifaceted Molecule, in Innate Immune Responses against Pathogens. Mediators of Inflammation, 2017, 9247574.

End, C., Bikker, F., Renner, M., Bergmann, G., Lyer, S., Blaich, S., Hudler, M., et al. (2009). DMBT1 functions as pattern-recognition molecule for poly-sulfated and poly-phosphorylated ligands. European Journal of Immunology, 39(3), 833–842.

Friedrich, M, Diegelmann, J., Schauber, J., Auernhammer, C. J., & Brand, S. (2015). Intestinal neuroendocrine cells and goblet cells are mediators of IL-17A-amplified epithelial IL-17C production in human inflammatory bowel disease. Mucosal Immunology, 8(4), 943–958.

Friedrich, Matthias, Pohin, M., Jackson, M. A., Korsunsky, I., Bullers, S. J., Rue-Albrecht, K., Christoforidou, Z., et al. (2021). IL-1-driven stromal-neutrophil interactions define a subset of patients with inflammatory bowel disease that does not respond to therapies. Nature Medicine, 27(11), 1970–1981.

Gonzalez-Acera, M., Patankar, J. V., Erkert, L., Cineus, R., Gamez-Belmonte, R., Leupold, T., Bubeck, M., et al. (2025). Integrated multimodel analysis of intestinal inflammation exposes key molecular features of preclinical and clinical IBD. Gut, 74(10), 1602–1615.

Hao, Y., Stuart, T., Kowalski, M. H., Choudhary, S., Hoffman, P., Hartman, A., Srivastava, A., et al. (2024). Dictionary learning for integrative, multimodal and scalable single-cell analysis. Nature Biotechnology, 42(2), 293–304.

Hracs, L., Windsor, J. W., Gorospe, J., Cummings, M., Coward, S., Buie, M. J., Quan, J., et al. (2025). Global evolution of inflammatory bowel disease across epidemiologic stages. Nature, 642(8067), 458–466.

Jansen, J. E., Aschenbrenner, D., Uhlig, H. H., Coles, M. C., & Gaffney, E. A. (2022). A method for the inference of cytokine interaction networks. PLoS Computational Biology, 18(6), e1010112.

Kaplan, G. G. (2025). The global burden of inflammatory bowel disease: from 2025 to 2045. Nature Reviews. Gastroenterology & Hepatology, 22(10), 708–720.

Kayal, M., Ungaro, R. C., Bader, G., Colombel, J.-F., Sandborn, W. J., & Stalgis, C. (2023). Net Remission Rates with Biologic Treatment in Crohn’s Disease: A Reappraisal of the Clinical Trial Data. Clinical Gastroenterology and Hepatology, 21(5), 1348–1350.

Korotkevich, G., Sukhov, V., Budin, N., Shpak, B., Artyomov, M. N., & Sergushichev, A. (2016). Fast gene set enrichment analysis. BioRxiv.

Linares, R., Gutiérrez, A., Márquez-Galera, Á., Caparrós, E., Aparicio, J. R., Madero, L., Payá, A., et al. (2022). Transcriptional regulation of chemokine network by biologic monotherapy in ileum of patients with Crohn’s disease. Biomedicine & Pharmacotherapy (Biomedecine & Pharmacotherapie), 147, 112653.

Marsal, J., Barreiro-de Acosta, M., Blumenstein, I., Cappello, M., Bazin, T., & Sebastian, S. (2022). Management of Non-response and Loss of Response to Anti-tumor Necrosis Factor Therapy in Inflammatory Bowel Disease. Frontiers in medicine, 9, 897936.

Mestas, J., & Hughes, C. C. W. (2004). Of mice and not men: differences between mouse and human immunology. Journal of Immunology, 172(5), 2731–2738.

Nies, J. F., & Panzer, U. (2020). IL-17C/IL-17RE: Emergence of a Unique Axis in TH17 Biology. Frontiers in Immunology, 11, 341.

Noor, N. M., Davies, N., Tahir, W., Bond, S., Dowling, F., Patel, K. V., Lyons, P. A., et al. (2025). Anti-tumor necrosis factor treatment from diagnosis is more effective and less costly than conventional “step-up” care for patients with active Crohn’s disease: a cost-effectiveness analysis from the PROFILE trial. Journal of Crohn’s & colitis, 19(9).

Olbei, M., Hautefort, I., Thomas, J. P., Csabai, L., Bohar, B., Koigi, S. S., Ibraheim, H., et al. (2026). Decoding cytokine networks in ulcerative colitis to identify pathogenic mechanisms and therapeutic targets. Science Signaling, 19(923), eadt0986.

Olbei, M., Thomas, J. P., Hautefort, I., Treveil, A., Bohar, B., Madgwick, M., Gul, L., et al. (2021). CytokineLink: A Cytokine Communication Map to Analyse Immune Responses-Case Studies in Inflammatory Bowel Disease and COVID-19. Cells, 10(9).

Qi, Z., Pang, W., Zha, X., Liu, Y., Liu, S., Xiao, F., Wang, X., et al. (2025). REG/Reg family proteins: mediating gut microbiota homeostasis and implications in digestive diseases. Gut microbes, 17(1), 2568055.

Schäfer, P. S. L., Dimitrov, D., Villablanca, E. J., & Saez-Rodriguez, J. (2024). Integrating single-cell multi-omics and prior biological knowledge for a functional characterization of the immune system. Nature Immunology, 25(3), 405–417.

Schmitt, H., Billmeier, U., Dieterich, W., Rath, T., Sonnewald, S., Reid, S., Hirschmann, S., et al. (2019). Expansion of IL-23 receptor bearing TNFR2+ T cells is associated with molecular resistance to anti-TNF therapy in Crohn’s disease. Gut, 68(5), 814–828.

Schmitt, H., Neurath, M. F., & Atreya, R. (2021). Role of the IL23/IL17 pathway in crohn’s disease. Frontiers in Immunology, 12, 622934.

Souza, R. F., Caetano, M. A. F., Magalhães, H. I. R., & Castelucci, P. (2023). Study of tumor necrosis factor receptor in the inflammatory bowel disease. World Journal of Gastroenterology, 29(18), 2733–2746.

Swedik, S., Madola, A., & Levine, A. (2021). IL-17C in human mucosal immunity: More than just a middle child. Cytokine, 146, 155641.

Thomas, T., Friedrich, M., Rich-Griffin, C., Pohin, M., Agarwal, D., Pakpoor, J., Lee, C., et al. (2024). A longitudinal single-cell atlas of anti-tumour necrosis factor treatment in inflammatory bowel disease. Nature Immunology, 25(11), 2152–2165.

Vebr, M., Pomahačová, R., Sýkora, J., & Schwarz, J. (2023). A narrative review of cytokine networks: pathophysiological and therapeutic implications for inflammatory bowel disease pathogenesis. Biomedicines, 11(12).

West, N. R., Hegazy, A. N., Owens, B. M. J., Bullers, S. J., Linggi, B., Buonocore, S., Coccia, M., et al. (2017). Oncostatin M drives intestinal inflammation and predicts response to tumor necrosis factor-neutralizing therapy in patients with inflammatory bowel disease. Nature Medicine, 23(5), 579–589.

