## Supplementary figures and images for "Cytokine interaction networks, not individual cytokines, drive anti-TNF response in Crohn’s disease"

### Fig_S1.png

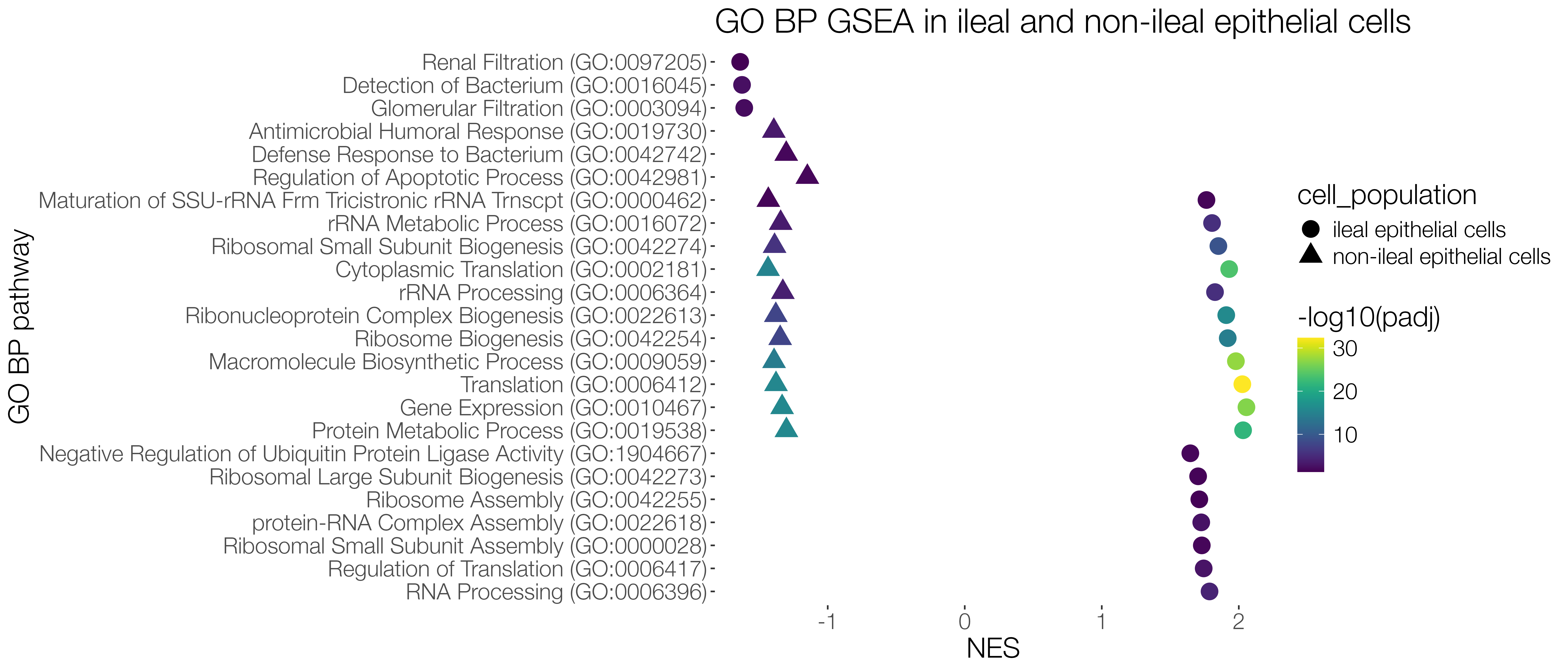

### Fig_S2.png

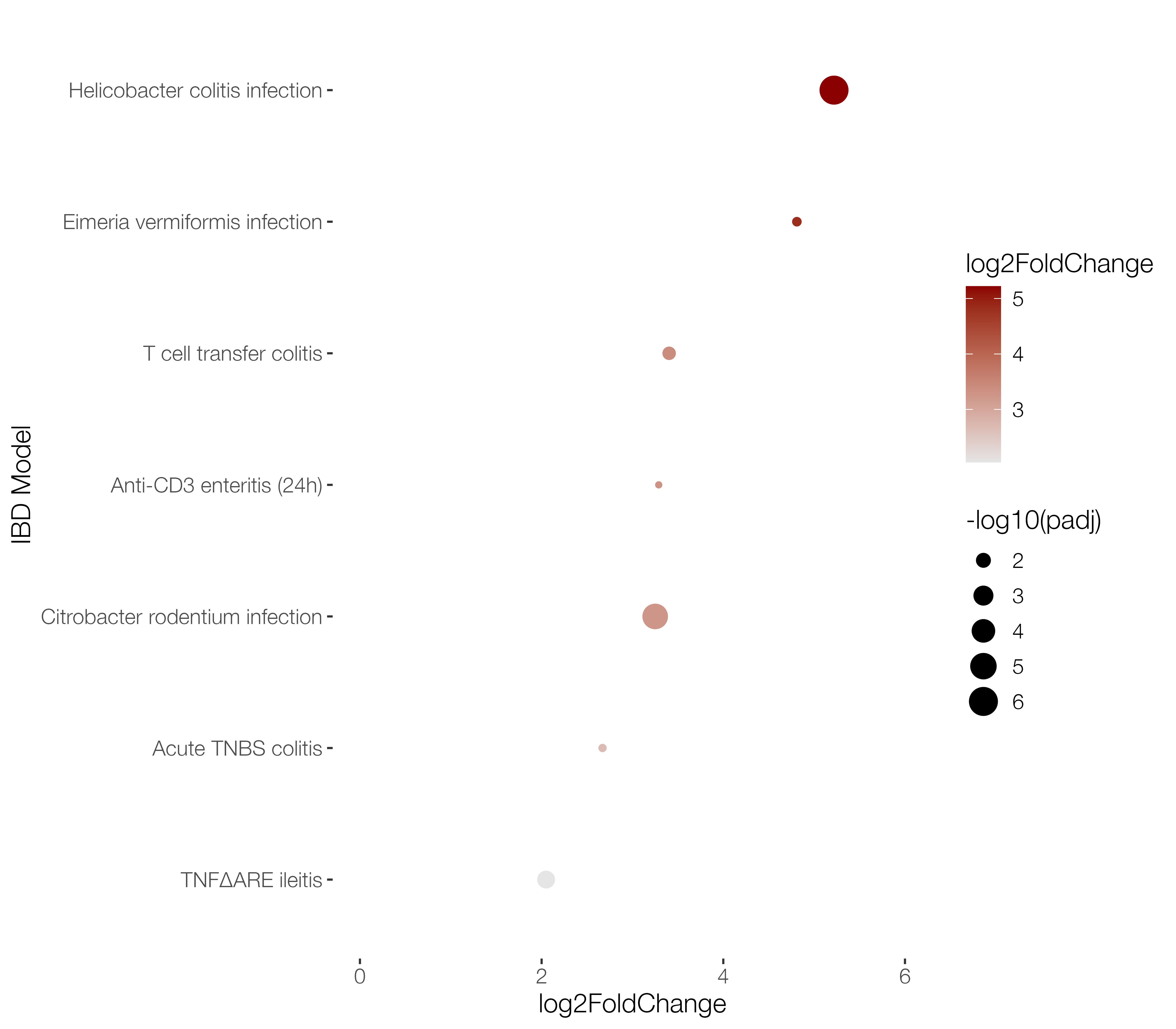

### Fig_S3.png

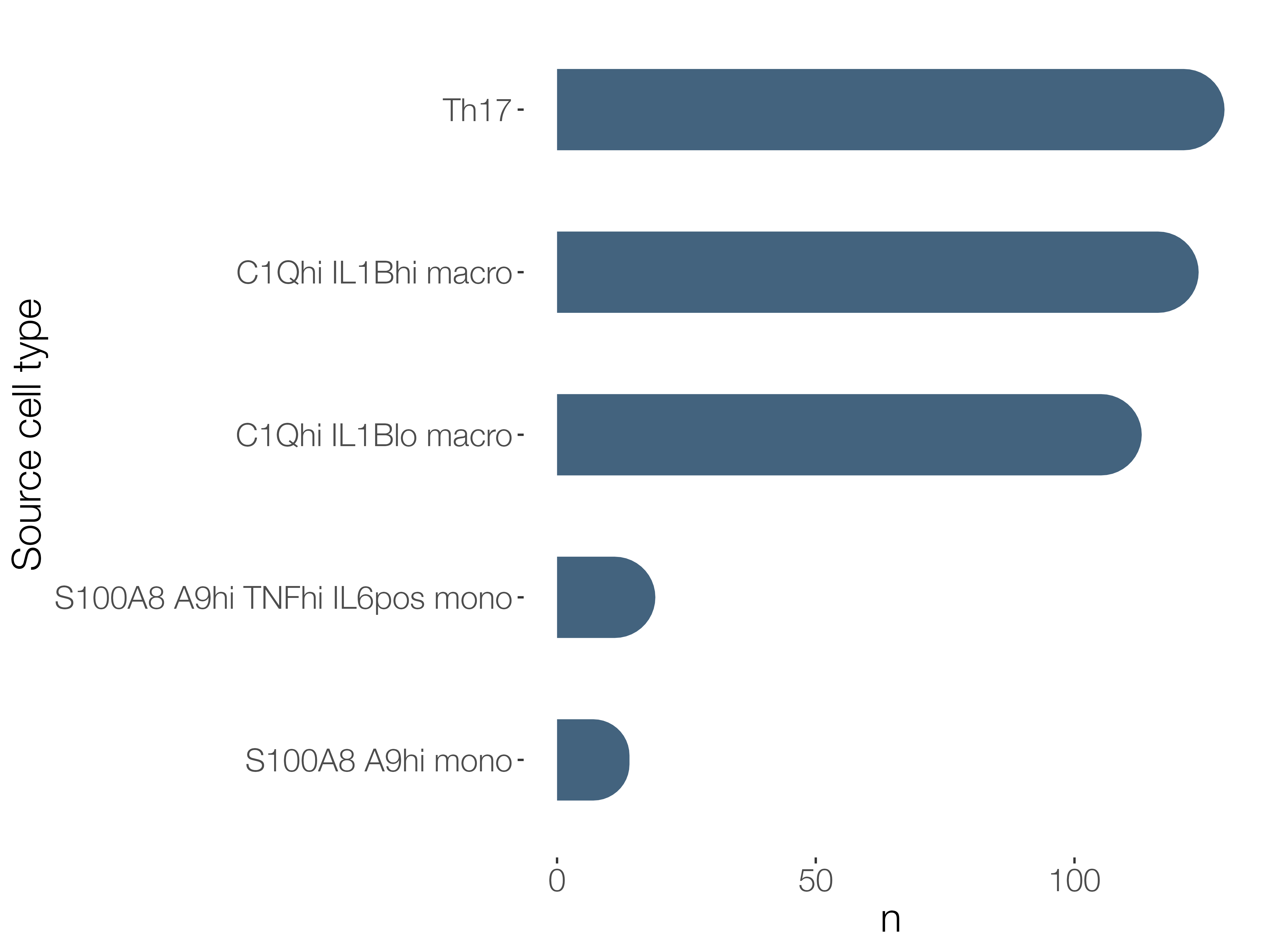
